# Concurrent Spatiotemporal Daily Land Use Regression Modeling and Missing Data Imputation of Fine Particulate Matter Using Distributed Space Time Expectation Maximization

**DOI:** 10.1101/354852

**Authors:** Seyed Mahmood Taghavi-Shahri, Alessandro Fassò, Behzad Mahaki, Heresh Amini

## Abstract

**Graphical Abstract:** 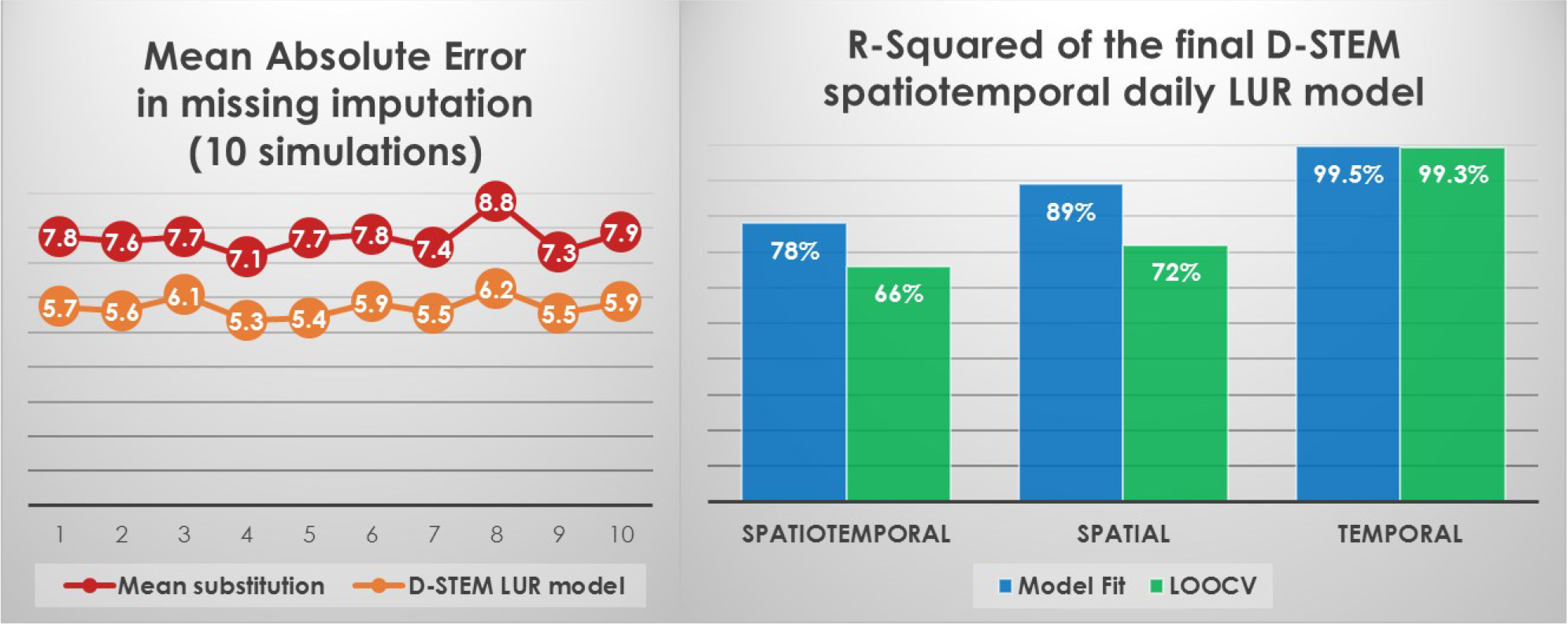

Land use regression (LUR) has been widely applied in epidemiologic research for exposure assessment. In this study, for the first time, we aimed to develop a spatiotemporal LUR model using Distributed Space Time Expectation Maximization (D-STEM). This spatiotemporal LUR model examined with daily particulate matter ≤ 2.5 μm (PM_2.5_) within the megacity of Tehran, capital of Iran. Moreover, D-STEM missing data imputation was compared with mean substitution in each monitoring station, as it is equivalent to ignoring of missing data, which is common in LUR studies that employ regulatory monitoring stations’ data. The amount of missing data was 28% of the total number of observations, in Tehran in 2015. The annual mean of PM_2.5_ concentrations was 33 μg/m^3^. Spatiotemporal R-squared of the D-STEM final daily LUR model was 78%, and leave-one-out cross-validation (LOOCV) R-squared was 66%. Spatial R-squared and LOOCV R-squared were 89% and 72%, respectively. Temporal R-squared and LOOCV R-squared were 99.5% and 99.3%, respectively. Mean absolute error decreased 26% in imputation of missing data by using the D-STEM final LUR model instead of mean substitution. This study reveals competence of the D-STEM software in spatiotemporal missing data imputation, estimation of temporal trend, and mapping of small scale (20 × 20 meters) within-city spatial variations, in the LUR context. The estimated PM_2.5_ concentrations maps could be used in future studies on short- and/or long-term health effects. Overall, we suggest using D-STEM capabilities in increasing LUR studies that employ data of regulatory network monitoring stations.

**Highlights:**

- First Land Use Regression using D-STEM, a recently introduced statistical software
- Assess D-STEM in spatiotemporal modeling, mapping, and missing data imputation
- Estimate high resolution (20×20 m) daily maps for exposure assessment in a megacity
- Provide both short- and long-term exposure assessment for epidemiological studies

## 1 Introduction

Air pollution caused 6.5 million deaths and 167 million disability-adjusted life-years (DALYs) in 2015 worldwide. In particular, ambient particulate matter with an aerodynamic diameter smaller than 2.5 μm (PM_2.5_), caused 4.2 million of deaths and 103 million DALYs worldwide in the same year (Forouzanfar et al., 2016).

Air pollution epidemiological studies need accurate exposure assessment and one frequently used method for individual exposure assessment is land use regression (LUR) modeling. LUR uses geographic independent variables (typically extracted using geographic information system (GIS)) as Potentially Predictive Variables (PPVs) to predict spatial variation of air pollution at unmonitored locations (Amini et al., 2017c; Hoek et al., 2008; Ryan and LeMasters, 2007).

Many LUR studies that based on data of regulatory network monitoring stations, use annual or seasonal mean of air pollutants with a subset of GIS generated PPVs in a multiple regression for spatial modeling and mapping (Huang et al., 2017; Kashima et al., 2018; Zou et al., 2015) that ignore temporal trend of air pollution. However, some other studies use relatively advanced models, such as linear mixed effect models, with satellite, meteorological, and land use predictors for spatiotemporal estimation of air pollution concentrations (Just et al., 2015; Kloog et al., 2011; Meng et al., 2016). However, the common issue is treating of air pollution missing data, and perhaps limited capability in accounting for temporal and spatial autocorrelations.

D-STEM (Distributed Space Time Expectation Maximization) is a comprehensive statistical software for analyzing and mapping of environmental data that was introduced in 2013, which is programmed and run in Matlab environment (The MathWorks). The underlying D-STEM general model accounts for spatial and temporal autocorrelations and measurement errors, spatial and temporal and spatiotemporal predictors, fixed and random effect coefficients, concurrent missing imputation and model fitting using Expectation Maximization (EM) algorithm. In addition, D-STEM supports multivariate modeling even in heterogonous stations (for example, simultaneous modeling of meteorological variables and air pollutants measured in the same or different stations), and data fusion capability for joint modeling (and calibration) of remote sensing data measured by a satellite along with air pollution concentrations measured in the ground stations (Finazzi, 2013; Finazzi and Fassò, 2014).

D-STEM software has been previously used in several air pollution studies in Europe (Calculli et al., 2015; Fassò, 2013; Fassò et al., 2016; Finazzi, 2013; Finazzi et al., 2013), but primary focus of those studies have been on statistical theory. Moreover, the resolution of their output maps has been about 1 km, which might be course for epidemiological studies that need individual-level exposure estimates. In addition, none of those studies used a pool of GIS generated PPVs for extensive independent variables selection. Overall, D-STEM has not been used so far for LUR modeling and for exposure estimation with very fine spatial resolution.

To date, several LUR studies have been published in Tehran to predict air pollution. These have been for particulate matter with an aerodynamic diameter smaller than 10 μm (PM_10_) and sulfur dioxide (SO_2_) (Amini et al., 2014), nitrogen oxides (NO, NO_2_, and NO_x_) (Amini et al., 2016), PM_2.5_, carbon monoxide (CO), and nitrogen dioxide (NO_2_) (Hassanpour Matikolaei et al., 2017), and alkylbenzenes (benzene, toluene, ethylbenzene, *m*-xylene, *p*-xylene, and *o*-xylene (BTEX), and total BTEX) (Amini et al., 2017b). In fact, multiple regression has been used in majority of these studies for modeling of annual or seasonal averages of air pollutants. However, the study of PM_2.5_, CO, and NO_2_ (Hassanpour Matikolaei et al., 2017), used panel regression to predict hourly concentrations with limited capability in accounting for spatial and temporal autocorrelations, and not supporting spatiotemporal missing imputation (Hassanpour Matikolaei et al., 2017).

In the present study, we aimed to utilize D-STEM with a relatively large pool of GIS generated PPVs for producing high resolution daily estimations of airborne PM_2.5_ in the Middle Eastern megacity of Tehran, Iran. We further aimed to assess some characteristics of the recently introduced D-STEM software. In addition, we aimed to compare performance of air pollution missing data imputation under two scenarios: (1) using mean substitution in each monitoring station, and (2) using D-STEM concurrent spatiotemporal missing imputation and LUR modeling.

## 2. Materials and Methods

### 2.1 Study area and data

The study area of this research is the megacity of Tehran, Iran which has 9 million urban residents and, due to diurnal migration, a daytime population of more than 10 million. Alborz Mountains are located in north of the city and there is a desert in the south. The populated area covers approximately 613 km^2^ and its elevation ranges from 1,000 to 1800 meters above sea level from south to north. The prevailing winds blow from west and north. The mean daily temperature ranges from about −15 °C in January to 43 °C in July with an annual average of 18.5 °C. The precipitation ranges from 0 millimeters (mm) in September to 40 mm in January, with an annual average of 150 mm. The weather of most days is sunny, and the mean cloud cover is about 30%. Geographic location of Tehran is in zone 39N of Universal Transverse Mercator (UTM) (Amini et al., 2016; Amini et al., 2014). Figure S1 of Supporting information shows the study area and location of monitoring stations.

Hourly data of fine particulate matter (PM_2.5_), which have been measured using fixed beta-gauge monitors, were obtained from Tehran Air Quality Control Company (AQCC) and Iranian Department of Environment (DOE) for year 2015. Daily PM_2.5_ was calculated by 24-hours averaging of hourly measured PM_2.5_ concentrations when days had complete records in all hours. Daily mean of PM_2.5_ for a set of 19 fixed measuring sites and for 365 days of year 2015 was used as response variable in this study.

A pool of 212 potentially predictive variables (PPVs) in six classes and 73 sub-classes were used for selecting model independent variables. These variables were in raster format for whole study area with resolution of 5×5 meters. The six GIS generated classes were Distance (60 variables), Product terms (52 variables), Land Use (50 variables), Traffic Surrogates (26 variables), Population Density (22 variables), and Geographic Location (2 variables). The full list of our PPVs could be found elsewhere (Amini et al., 2016; Amini et al., 2014).

### 2.2 Model Specification

D-STEM spatiotemporal LUR model for PM_2.5_ is given by the following equation where Y is response variable, which is spatiotemporally indexed at location s and time t, provided X’s the GIS generated spatial predictors that are assumed temporally constant over time and spatially indexed at location s:

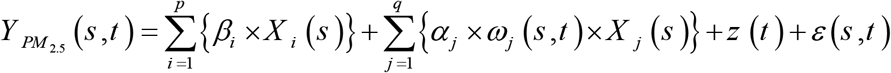

While the left-hand side of the above equation defines the response variable, the right-hand side of this equation consists of four parts. First, sum of p summand: *β*_*i*_ multiplied by *X*_*i*_ (*s*), which are a fixed effect coefficient and the corresponding GIS generated spatial predictor. Second, sum of q summand: (*α*_*j*_ multiplied by *ω*_*j*_ (*s*,*t*) and multiplied by *X*_*j*_ (*s*,*t*), which are a scale changing coefficient and the corresponding spatiotemporal random effect coefficient, and the corresponding GIS generated spatial predictor, respectively. Third part is *z* (*t*) which is a latent temporal state variable that concurrently estimated using a sub-model. Forth part is *ε*(*s*,*t*) which denotes measurement error that assumed normally distributed with zero mean and uncorrelated over space and time.

Latent spatiotemporal random effect coefficients *ω*_*j*_ (*s*,*t*)assumed to be independent of each other and normally distributed with zero mean and unit variance that uncorrelated over time but correlated over space with exponential spatial auto-covariance function 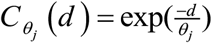 where d is the distance between two generic spatial points and *θ*_*j*_ is auto-covariance parameter.

Since the latent spatiotemporal random effect coefficients (*ω*_*j*_ (*s*,*t*)assumed to have zero means, we are not allowed to incorporate any GIS generated spatial predictor in random effect part, except if we already used it in fixed effect part. In the current representation, *β* could be interpreted as global effect of GIS generated predictor, and *α* could be used for testing spatiotemporal variation in the effect of corresponding predictor.

Finally, latent temporal state variable (third part of model) estimated concurrently using following sub-model:

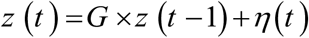

Where G is coefficient of temporal state transition and *η* is temporal state innovation that assumed normally distributed with zero mean.

The estimated parameters of the D-STEM spatiotemporal LUR model are as follows: fixed effect coefficients, scaling changing coefficients of the corresponding spatiotemporal random effect coefficients, parameters of the underlying auto-covariance functions of the corresponding spatiotemporal random effect coefficients, temporal state transition coefficient, variance of the temporal state innovation, and variance of the measurement error.

### 2.3 Model fitting, assessing, and mapping

A forward algorithm, based on maximization of spatial Leave-One-Out Cross-Validation (LOOCV) R-squared and consistency with prior knowledge, was used to select suitable predictors from pool of PPVs. This algorithm considers sign consistency of fixed effect coefficients with a priori assumed direction of effects. Effect directions defined as positive, negative, or arbitrary (unknown) based on knowledge about emission of air pollutions.

The final daily LUR model was assessed in three forms: a) spatiotemporal, b) spatial (temporally averaged to assess spatial variations), and c) temporal (spatially averaged to assess temporal variations). Also, Normalized Mean Bias Factor (NMBF) was used for checking whether models underestimate or overestimate observed data (Yu et al., 2006).

The power of missing data imputation was compared between mean substitution in each monitoring station, and final spatiotemporal LUR model by 10 independent simulations. In each simulation 10% of spatiotemporal observed data was randomly selected and was replaced by missing data. Afterwards, both models were independently employed for missing imputation. The predicted values of the randomly missed data and original observed values (that randomly missed), was used to compute Mean Absolute Error (MAE) in missing imputation of each model. Finally, MAEs that computed in these independent simulations and percent of MAEs change between two models, was averaged and reported.

Mapping was done using D-STEM for each day, by providing GIS generated maps that correspond to predictors of the final daily LUR model. Seasonal maps were generated by averaging of daily output maps in the corresponding time periods. Finally, we applied a quantification limit (QL) that its lower bound was the minimum of within monitoring station averages of observed concentration in the corresponding time period divided by square root of two, and its upper bounds was 120% of the maximum of within monitoring station averages of observed concentration in the same time period. Grid cells in the final output maps that had values outside QL interval were set to nearest QL bound and frequency of these corrections was calculated and reported (Amini et al., 2016; Amini et al., 2014; Henderson et al., 2007)

## 3. Results and Discussion

### 3.1 Description of data

Number of PM_2.5_ daily observations at 19 monitoring stations was 5016. Frequency of missing data varies from 12% to 47% in monitoring stations with an average of 28% over all stations. Frequency of available data for each day varied from 2 stations to 19 stations with an average of 14 stations. Data availability pattern of daily PM_2.5_ in monitoring stations of Tehran in 2015 is presented in Figure S2 of Supporting information.

Annual mean of observed PM_2.5_ concentrations (spatially and temporally average of monitoring stations) was 33 μg/m^3^ that is more than three times of World Health Organization (WHO) guideline (10 μg/m^3^) and about three times of US Environmental Protection Agency (EPA) guideline (12 μg/m^3^) (US EPA, 2013; WHO, 2006), with a range of 21 μg/m^3^ to 48 μg/m^3^ in different monitoring stations. This is in line with the findings of Brauer et al. (2016) and Shaddick et al. (2018) where they reported very high population-weighted exposure estimates for PM_2.5_ in the Middle East (Brauer et al., 2016; Shaddick et al., 2018). Faridi et al. (2018) also reported very similar annual mean for PM_2.5_ in Tehran (Faridi et al., 2018). Daily mean of Tehran PM_2.5_ concentrations in 2015 (spatially average of monitoring stations) was greater than WHO daily standard (25 μg/m^3^) in 246 days (67% of year), and was greater than US EPA daily standard (35 μg/m^3^) in 121 days (33% of year), and eventually was greater than 55 μg/m^3^ in 31 days. As a result, the air quality index (AQI) in Tehran in 2015 was unhealthy for at least sensitive groups in one third of year, and was unhealthy for all groups in about one month (8.6% of year that exceeded 55 μg/m^3^), based on US EPA breakpoints (US EPA, 2013; WHO, 2006). The very high values of air pollution have been confirmed in large Tehran Study of Exposure Prediction for Environmental Health Research (SEPEHR) measurement campaigns (Amini et al., 2017a).

### 3.2 Description of the LUR model

Natural logarithm of PM_2.5_ was standardized (using mean and standard deviation of its all spatiotemporal values) and utilized in spatiotemporal LUR model as response variable. Also, standardized version of GIS generated PPVs was employed in the model. Summary statistics for these variables are presented in Table S1 of Supporting information.

Table 1 presents estimated parameters of spatiotemporal daily LUR model. This model accounts for both temporal and spatial autocorrelations, and moreover supports inclusion of predictors for concurrent missing imputation and LUR model fitting. Among 212 GIS generated predictors that produced a huge possible combination of predictors, eventually 8 predictors were selected by our described algorithm to include in the spatiotemporal final daily LUR model.

**Table 1.**
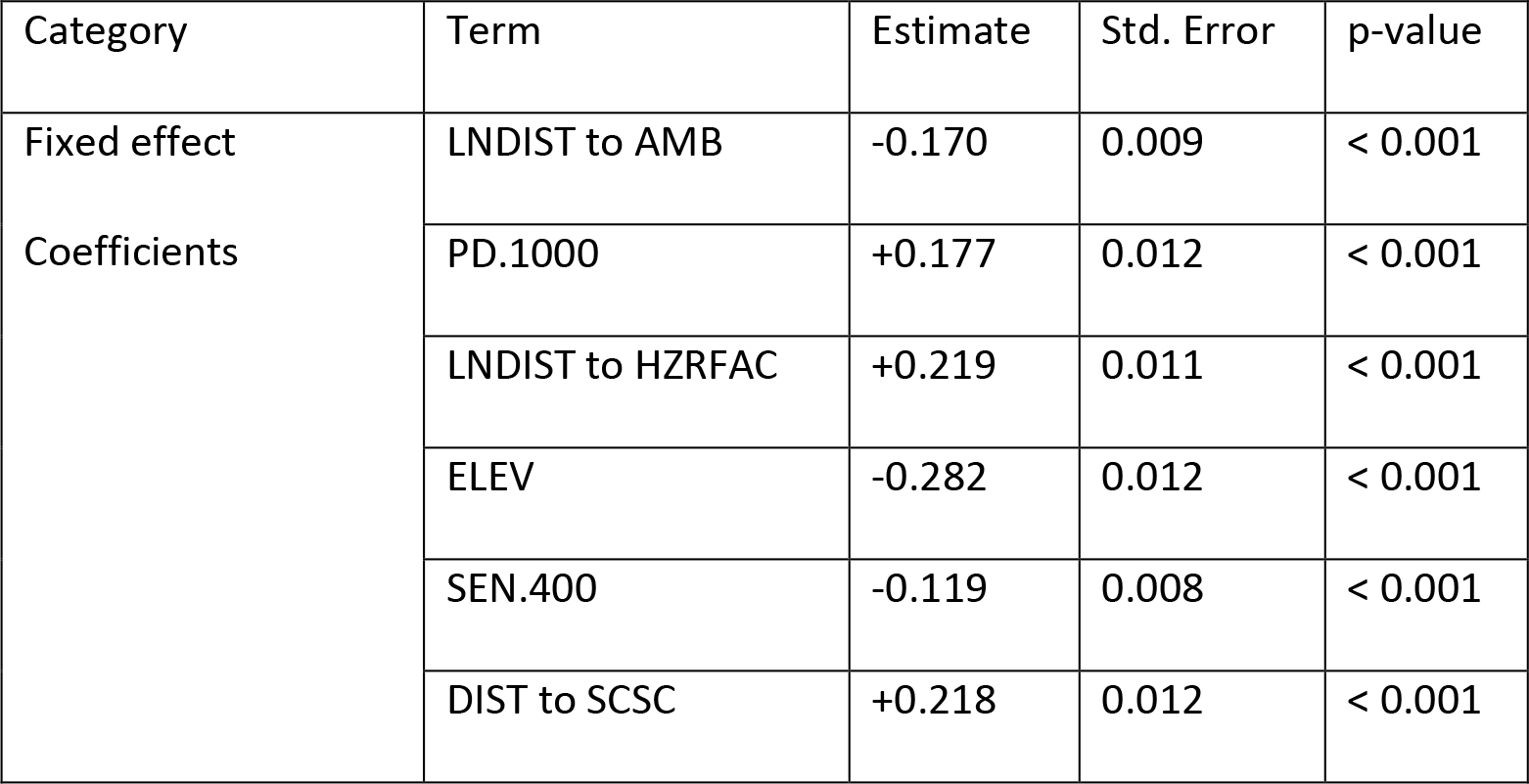
Estimated parameters of spatiotemporal final daily LUR model for PM_2.5_ in Tehran 2015.

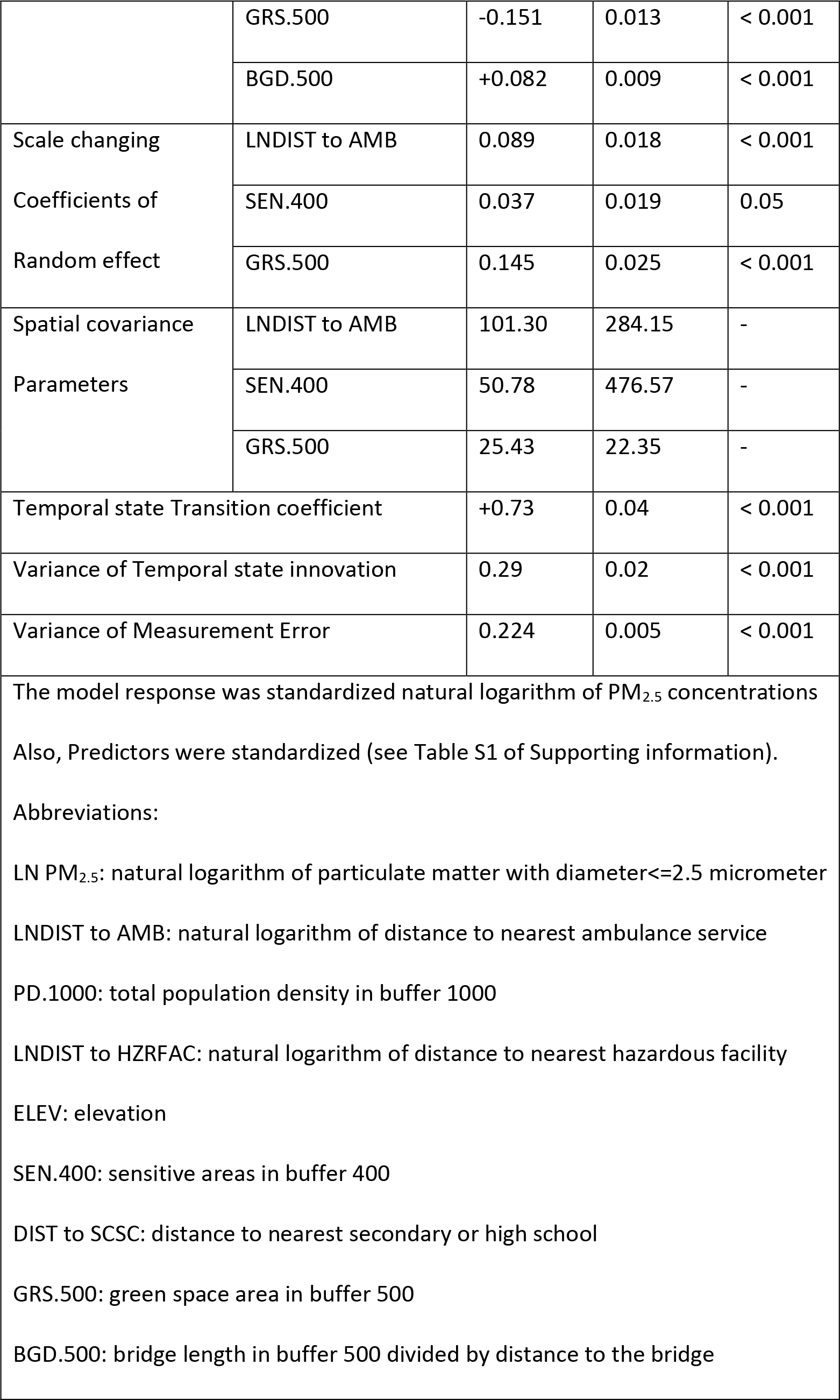

GIS generated predictors that had a positive coefficient in the final spatiotemporal daily LUR model were: total population density in buffer of 1000 m, natural logarithm of distance to nearest hazardous facility (with an arbitrary assumed direction of the effect), distance to the nearest secondary or high school, and bridge length in buffer of 500 m divided by distance to the bridges.

On the other hand, the predictors that had a negative coefficient in the final spatiotemporal daily LUR model were: natural logarithm of distance to nearest ambulance service (assumed sign was arbitrary), elevation (assumed sign was arbitrary), sensitive areas in buffer of 400 m (assumed sign was arbitrary), and green space areas in buffer of 500 m.

Random effect predictors in the final spatiotemporal daily LUR model were as follow: natural logarithm of distance to nearest ambulance service, sensitive areas in buffer of 400 m, and green space areas in buffer of 500 m. The corresponding scale changing coefficients were significant or borderline significant. That means coefficient of these three predictors vary in time and space, and observed data are spatially auto-correlated.

Temporal state transition coefficient was significant (p< 0.001) and positive that indicates a positive temporal autocorrelation existed between successive observations. Moreover, variance of the temporal state innovation was significant (p< 0.001) that means variations existed in temporal trend of air pollution.

### 3.3 Assessing fit of the LUR model

Figure 1 shows temporal pattern (spatially averaged) of daily PM_2.5_ for observed and predicted data of the final daily LUR model. Figure 1 confirms that D-STEM succeeds in capturing natural turbulence of PM_2.5_ time series, even without employing any temporal predictors.

**Figure 1.**
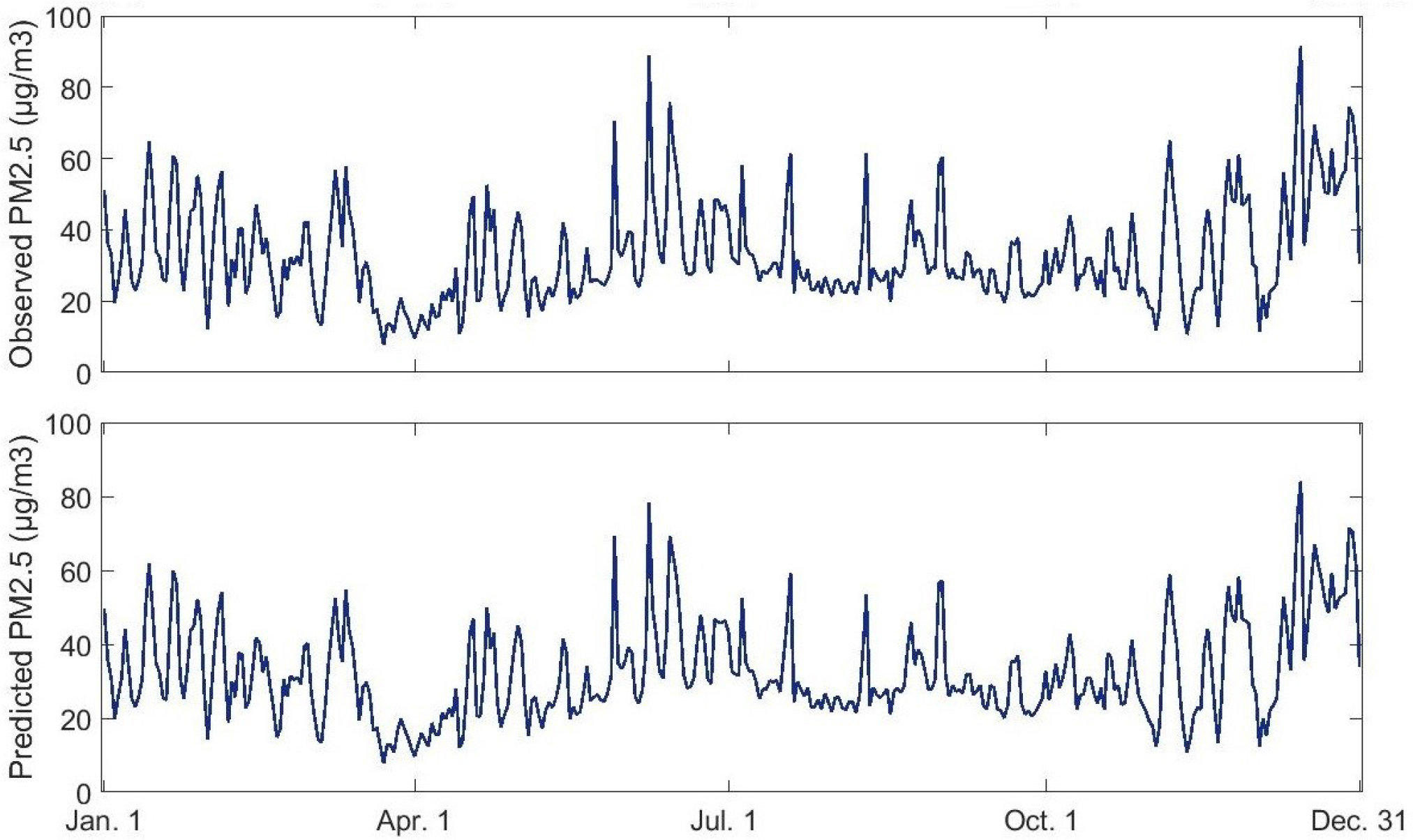
Temporal pattern (spatially averaged) of daily PM_2.5_ in Tehran 2015 (observed and predicted).

Spatiotemporal R-squared and spatiotemporal LOOCV R-squared were 78% and 66%, respectively. Spatial R-squared (temporally averaged) and spatial LOOCV R-squared were 89% and 72%, respectively. Temporal R-squared (spatially averaged) and temporal LOOCV R-squared were 99.5% and 99.3%, respectively. Root Mean Square Error (RMSE) criteria are shown in Table S2 of Supporting information for further assessment.

NMBF of the spatiotemporal final daily LUR model was −0.03 that means on average this model slightly underestimated observed data (with a factor of less than 3 percent). This slight underestimation also confirmed in Figure 1 and may be due to some large positive outliers that were not handled appropriately. Unfortunately, this criterion has not been reported in other studies that applied D-STEM (Calculli et al., 2015; Fassò, 2013; Fassò et al., 2016; Finazzi, 2013; Finazzi et al., 2013), so it is not clear that this slight deficiency is belonging to D-STEM software or it is an accidental observation that happened in our study.

However, the temporal prediction of D-STEM LUR model was very well. In fact, temporal R-squared and temporal LOOCV R-squared were greater than 99%. In a study by Kloog et al. (Kloog et al., 2011) where they employed land use and satellite-derived Aerosol Optical Depth (AOD) and multiple stations (spatiotemporal) meteorological predictors in a mixed model, the temporal cross-validated R-squared was 84%. Note that we did not use any spatiotemporal or pure temporal predictors in the D-STEM software. A similar study in Tehran that modelled PM_2.5_ with a panel regression used pure temporal predictors, i.e., one station meteorological variables, just in order to partly capture temporal innovations of air pollution data (Hassanpour Matikolaei et al., 2017). Hence, the use of pure temporal predictors in D-STEM for spatiotemporal modeling of air pollution is completely unnecessary.

The spatial R-squared of our D-STEM final LUR model (and even spatial LOOCV R-squared) was greater than 65% that reported in a recently published study on LUR modelling of PM_2.5_ in Tehran. That study, that used a panel regression for spatiotemporal modeling of fixed monitoring air pollution data, provided GIS generated spatial data, meteorological data of one station, and real time google traffic data as predictors (Hassanpour Matikolaei et al., 2017). Therefore, despite of the fact that predictors of our D-STEM spatiotemporal LUR model were limited to GIS generated spatial data, D-STEM performed better.

### 3.4 Assessing missing data imputation

Mean Absolute error (MAE) in 10 independent simulations for spatiotemporal missing imputation, was 5.7 μg/m^3^ with a range of 5.3 μg/m^3^ to 6.2 μg/m^3^ in the D-STEM final spatiotemporal daily LUR model, and was 7.7 μg/m^3^ with a range of 7.1 μg/m^3^ to 8.8 μg/m^3^ in Mean substitution in each monitoring station.

Changes between MAEs showed a decreased range from 24.3% to 29.9% with an average of 25.9% in using D-STEM final spatiotemporal daily LUR model instead of Mean substitution in each monitoring station. Details of MAEs and changes between MAEs are presented in Table S3 of Supporting information.

This result is important because missing data exists in almost any air pollution study of fixed air monitoring stations, such as our two previous LUR studies of PM_10_, SO_2_, and nitrogen oxides in Tehran. In those studies we imputed air pollution missing data using EM algorithm in a multivariate way but without considering spatial and temporal autocorrelations (Amini et al., 2016; Amini et al., 2014). Although, some other LUR studies such as one earlier mentioned LUR study of PM_2.5_ in Tehran that modeled using a panel regression, did not describe the missing data issue (Hassanpour Matikolaei et al., 2017).

This study proved that D-STEM missing imputation is better than ignoring of missing data, which is common at least when the amount of missing data is small (Meng et al., 2016). We believe D-STEM is worthy to be applied in missing imputation of (fixed monitoring stations) air pollution data, because it imputes missing data concurrently with modeling, and we see that the underlying temporal model of D-STEM perform very well. Moreover, D-STEM can capture spatial variation of air pollution data by incorporating suitable land use and GIS generated predictors. Furthermore, D-STEM spatiotemporal model that concurrently impute missing data has ability to employ spatiotemporal meteorological predictors, and even remote sensing data.

### 3.5 Generating output maps

We produced daily PM_2.5_ concentrations maps with resolution of 20×20 meters for whole study area that cover 613 km^2^. Cooler and warmer season maps of PM_2.5_ were produced by averaging of D-STEM generated daily maps in the corresponding periods.

Quantification limits (QL) that calculated based on averaged observed values of air quality monitoring stations in the corresponding periods, had a lower bound of 13.6 μg/m^3^ in cooler season, and 15.2 μg/m^3^ in warmer season; and had upper bound of 68.1 μg/m^3^ in cooler season, and 50.1 μg/m^3^ in warmer season.

Values beyond these quantification limits were corrected by setting to the nearest upper or lower limits. Percent of these corrections for lower and upper bounds of the corresponding QL limits were 0.03% and 1.9% in the cooler season map, and were 0.1% and 3.4% in warmer season map, respectively. Hence, the under- or over-estimation of PM_2.5_ in the D-STEM LUR generated maps were relatively small when compared to other LUR studies (Amini et al., 2016; Amini et al., 2014).

The seasonal estimation maps of PM_2.5_ concentrations in Tehran in 2015, with resolution of 20×20 meters, are presented in Figure 2. The scatter plot of predicted versus observed values are presented in Figure S3 of Supporting information for cooler and warmer seasons maps of PM_2.5_ concentrations.

**Figure 2.**
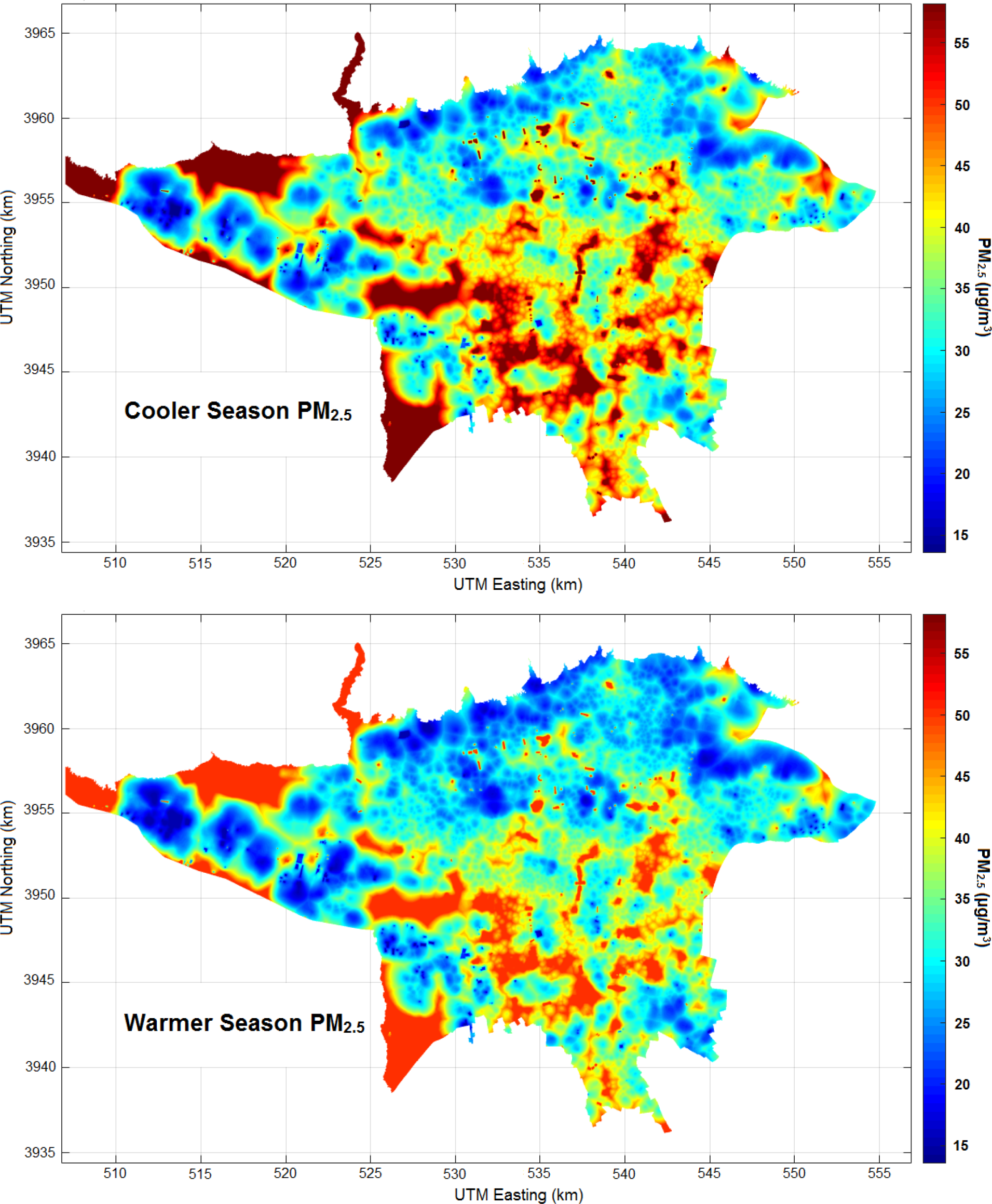
Estimated cooler and warmer seasons maps of PM_2.5_ in Tehran 2015 with resolution of 20×20 meters that produced by averaging of daily LUR maps over corresponding periods.

## 4. Conclusions

This was first LUR study that used recently introduced comprehensive statistical software of D-STEM. We used D-STEM in spatiotemporal missing imputation, modeling and mapping of airborne PM_2.5_ in Middle Eastern megacity of Tehran, where concentrations of PM_2.5_ was more than three times of WHO guideline. The estimated maps using D-STEM daily PM_2.5_ LUR model have resolution of 20×20 meters (for whole study area that covers 613 km^2^), while previous studies that employed D-STEM (in Europe), produced estimation maps with resolution of about 1 km (and all of them were in non-LUR context).

Our study demonstrated that D-STEM performed very well in temporal modeling of PM_2.5_ during year 2015 in Tehran, even without using any temporal or spatiotemporal predictors. Moreover, D-STEM daily spatiotemporal and spatial prediction and cross-validated prediction were quite good. This could benefit future epidemiological studies with both short-term and long-term exposure estimates. Furthermore, D-STEM build-in missing imputation capability compared with mean substitution in each monitoring station, which is equivalent to ignoring of missing data, proves usefulness of D-STEM LUR model in missing imputation of air pollution data.

Nonetheless, further research is needed to explore more advance features of D-STEM software, such as joint modeling of several air pollutants, joint modeling of one or several air pollutants with meteorological variables, and incorporation of remotely sensed data. Overall, we suggest using D-STEM capabilities in increasing LUR studies that employ data of regulatory network monitoring stations.

## Acknowledgements

This research was part of the first author’s Ph.D. thesis in Biostatistics and was supported by Isfahan University of Medical Sciences [grant number 394937]. The authors acknowledge Tehran Air Quality Control Company and Iranian Department of Environment for providing air pollution data.

## Declaration of interest

None.

## Appendix A. Supporting Information

Figures: Study area and geographical location of air quality monitoring stations, data availability pattern of observed daily PM_2.5_ in each monitoring station, scatter plot of predicted versus observed values for cooler and warmer seasons maps of PM_2.5_ concentrations.

Tables: descriptive statistics for response and predictors, Assessment and validation metrics, Mean Absolute Error (MAE) in 10 independent simulations, D-STEM output of the final model.

